# Irradiation of UVC LED at 277 nm inactivates coronaviruses by photodegradation of spike protein

**DOI:** 10.1101/2021.05.31.446403

**Authors:** Qunxiang Ong, J. W. Ronnie Teo, Joshua Dela Cruz, Elijah Wee, Winson Wee, Weiping Han

## Abstract

To interrupt SARS-CoV-2 transmission chains, Ultraviolet-C (UVC) irradiation has emerged as a potential disinfection tool to aid in blocking the spread of coronaviruses. While conventional 254-nm UVC mercury lamps have been used for disinfection purposes, other UVC wavelengths have emerged as attractive alternatives but a direct comparison of these tools is lacking with the inherent mechanistic properties unclear. Our results using human coronaviruses, hCoV-229E and hCoV-OC43, have indicated that 277-nm UVC LED is most effective in viral inactivation, followed by 222-nm far UVC and 254-nm UVC mercury lamp. While UVC mercury lamp is more effective in degrading viral genomic content compared to 277-nm UVC LED, the latter results in a pronounced photo-degradation of spike proteins which potentially contributed to the higher efficacy of coronavirus inactivation. Hence, inactivation of coronaviruses by 277-nm UVC LED irradiation constitutes a more promising method for disinfection.

## INTRODUCTION

The novel coronavirus SARS-CoV-2 has precipitated into the COVID-19 pandemic, and at the time of writing, resulted in more than 171 million infections and 3 million deaths. The actual numbers should be much higher than reported, given the high incidence of asymptomatic cases escaping the capture by traditional diagnostic methods. Vaccination, masking, rigorous testing and thorough public disinfection strategies become vital prongs in combating virus spread within communities. Amongst the latter, ultraviolet irradiation presents as an attractive strategy, given its use being well established in inactivating viruses and killing other microbes(Lin et al., 2020; Raeiszadeh and Adeli, 2020). Consequently, UVC mercury lamps have been increasingly deployed in hospital settings.

UVC has been well known to possess germicidal properties and inactivate pathogenic microbes by damaging nucleic acids and proteins, thereby eliminating their ability to reproduce(Rauth, 1965; Sehgal, 1973; Setlow and Carrier, 1966; Werbin et al., 1966). The mechanism at which UV inactivates microbes depends highly on the specific wavelengths. 277nm UVC LEDs (Beck et al., 2017; Kim and Kang, 2018; Nguyen et al., 2019; Nunayon et al., 2020) and far UVC sources(Welch et al., 2018a, 2018b) have recently emerged as attractive alternatives to UVC mercury lamps. The former does not require mercury, which is banned by the Minamata Convention, has very short turn-on time, and is generally more reliable and has a longer lifetime(Song et al., 2016). The far UVC, on the other hand, is shown to be effective in inactivation of bacteria and human coronaviruses, and potentially poses less safety concerns for deployment(Barnard et al., 2020; Buonanno et al., 2020).

Numerous studies have studied the sensitivity of different microbes to UVC wavelengths, including human coronaviruses (Buonanno et al., 2020; Gerchman et al., 2020)and SARS-CoV-2(Inagaki et al., 2020; Kitagawa et al., 2020; Storm et al., 2020). However, no study to date has performed a direct comparative study on the efficacy of different UVC wavelengths on inactivation of coronaviruses. In addition, mechanistic insight into how different UVC wavelengths inactivate coronaviruses is severely lacking, and greater understanding in this area would facilitate their deployment in future pandemics. Here, we utilized human coronaviruses, HCoV-229E and OC43, for efficacy studies where 277nm UVC LED consistently outperforms the other UVC wavelengths in inactivating coronaviruses. Mechanistic studies suggest that this is achieved via a combination of photo-degradation of spike proteins and RNA molecules.

## RESULTS

### Utilizing human coronaviruses for UVC-induced inactivation studies

Coronaviruses are split into the four genera: Alphacoronavirus, Betacoronavirus, Gammacoronavirus and Deltacoronavirus(Fehr and Perlman, 2015; Woo et al., 2010). Among the different genera, alphacoronaviruses and betacoronaviruses have been known to infect mammals, and pose as a significant risk to the human population. The betacoronaviruses, MERS-CoV, SARS-CoV and SARS-CoV-2, may produce severe symptoms in patients while hCoV-OC43 and hCoV-229E cause about 15% of common colds (**Fig. 1A**). In terms of genomic organization, coronaviruses are the largest enveloped RNA viruses with positive single-stranded RNA molecules from 27 to 32 kilobases. The genome comprises of the replicase gene that encodes for the non-structural proteins of the genomes at about 20 kilobases, while similar structural proteins in the form of spike, envelop, membrane and nucleocapsid proteins are interspersed at the 3’ end of the genome (**Fig. 1B**).

**Figure 1.**
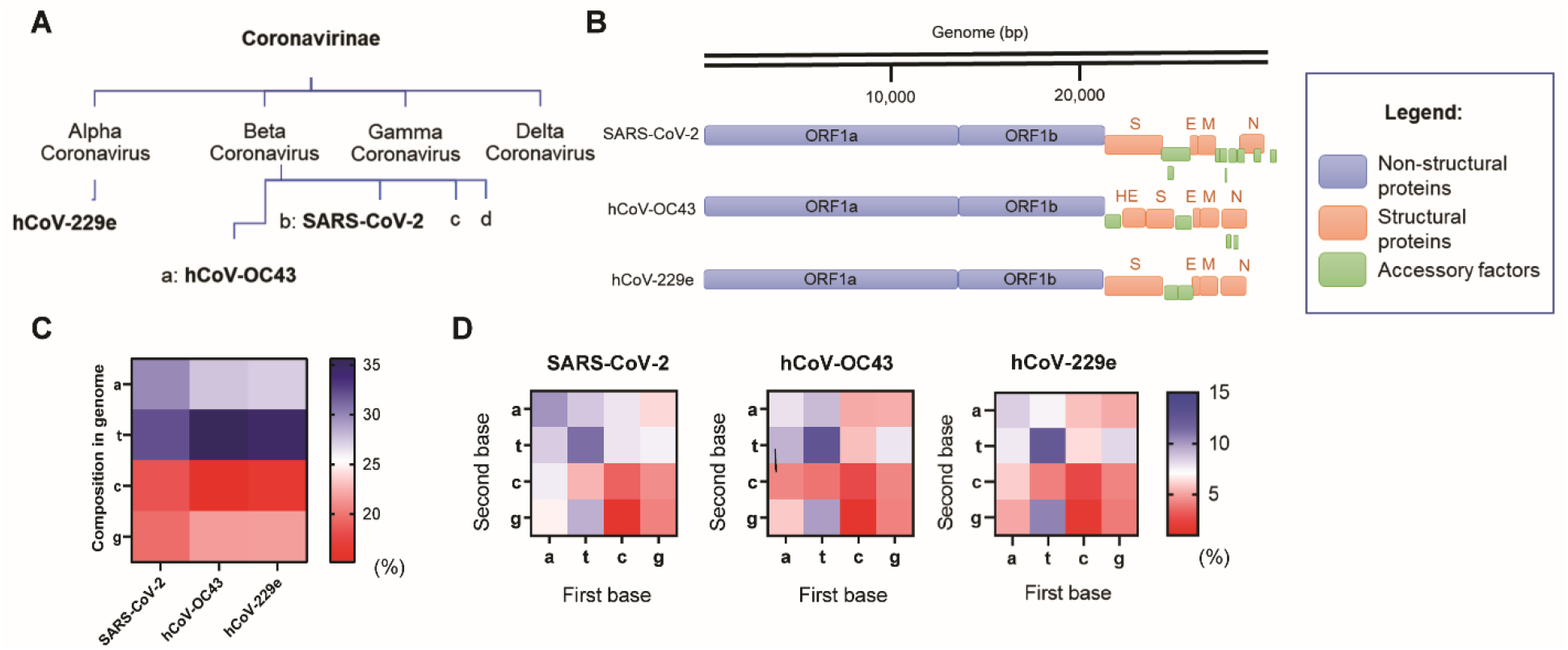
Utilizing human coronaviruses for UVC-induced inactivation studies. (a) The different genera of the coronavirus family. Alpha and beta-coronaviruses with the various highlighted viruses, hCoV-229e, hCoV-OC43 and SARS-CoV-2. (b) Genome organizations of SARS-CoV-2, hCoV-OC43 and hCoV-229e. (c) Overall base composition of SARS-CoV-2, hCoV-OC43 and hCoV-229e. (d) Adjacent base composition of SARS-CoV-2, hCoV-OC43 and hCoV-229e.

It has been established that RNA chains are directly disrupted by UVC by formation of pyrimidine dimers(Merriam and Gordon, 1967). The dimerization reaction occurs from adjacent pyrimidine bases in the form of uracil and cytosine(Beukers et al., 2010; Brown and Johns, 1968). We analyzed the genomic content of the coronaviruses based on these sequences as indicated their accession numbers from the NCBI Nucleotide: hCoV-OC43 (MW532119.1), hCoV-229e (KU291448.1) and SARS-CoV-2 (MW403500.1). We found that their overall base composition (**Fig. 1C**) and adjacent base arrangements (**Fig. 1D**) to be similar. We therefore hypothesize that the kinetics of UVC inactivation of hCoV-OC43 and hCoV-229E to be roughly similar.

### Inactivation of human coronaviruses after exposure to different UVC wavelengths

To examine the inactivation efficacy of UVC on hCoV-OC43 and hCoV-229E, virus was placed on plastic petri dishes and expose to various UVC wavelengths of 73 μW/cm^2^ for different timings ranging from 30 to 300 seconds (**Fig. S1-S2**). The reduction in infectivity of hCoV-OC43 (**Fig. 2A, Table S1**) and hCoV-229E (**Fig. 2B, Table S2**) can be observed after exposure to different UVC irradiation. The 277-nm UVC LED was most effective in carrying out the inactivation and achieved 3-log inactivation at 22 mJ/cm^2^ for both human coronavirus strains, whereas 254-nm UV lamp achieved only 2-log inactivation for hCoV-OC43 and 1-log inactivation for hCoV-229E with the same dosage. Two-way ANOVA analyses of both sets of data reveal significant differences (**Table S1-2**) between the three UVC wavelengths in carrying out coronavirus inactivation (p=0.0001 and p<0.0001 for hCoV-OC43 and hCoV-229E respectively).

**Figure 2.**
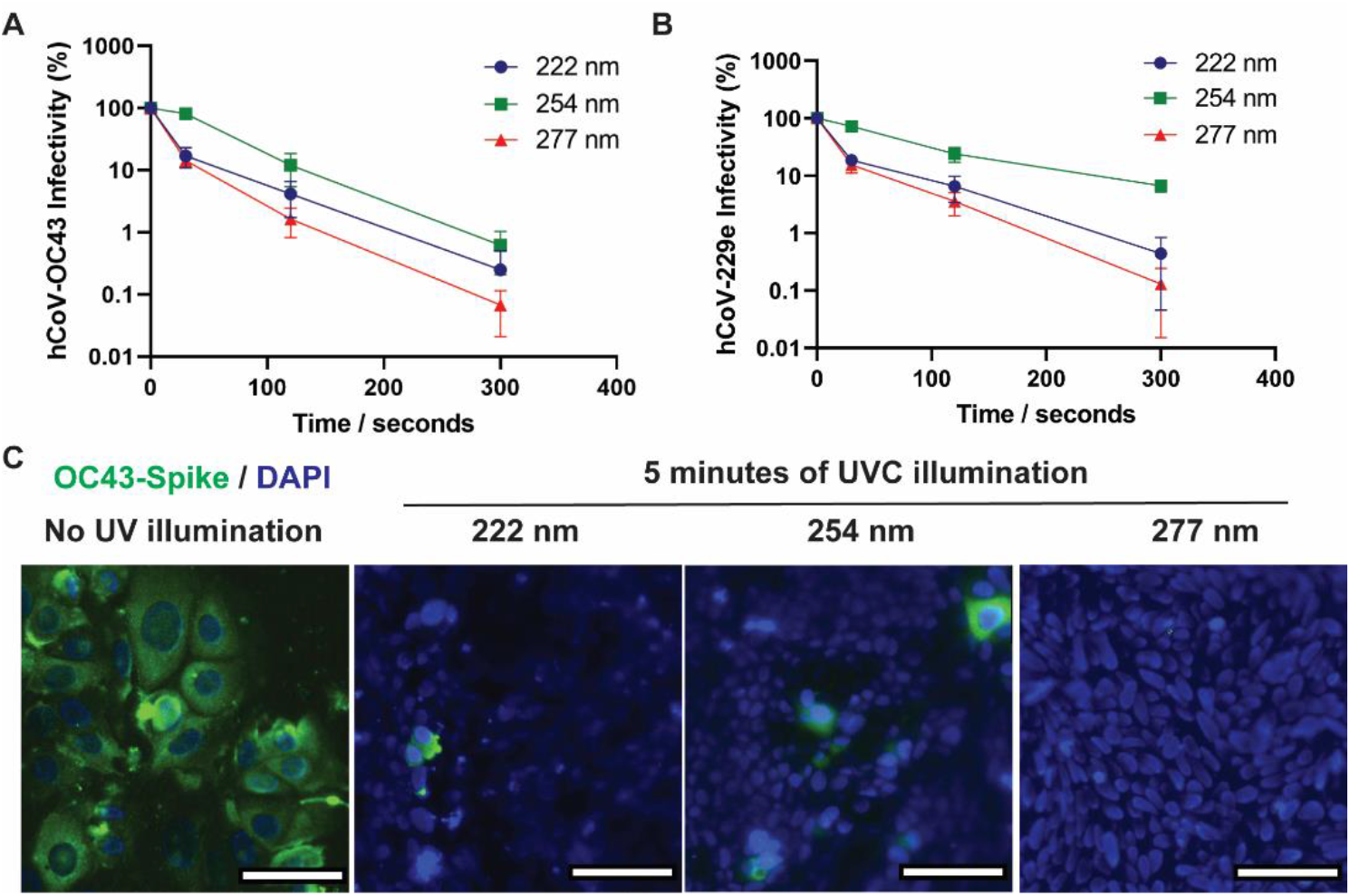
Inactivation of human coronaviruses after exposure to different UVC wavelengths. (a) HCoV-OC43 infectivity as a function of the duration of different UVC sources at 73 μW/cm^2^. Infectivity is defined as a function of PFU_UV_/PFU_NoUV_. Values are reported as mean +/− SD from n = 3 experiments. (b) HCoV-229e infectivity as a function of the duration of different UVC sources at 73 μW/cm^2^. Infectivity is defined as a function of PFU_UV_/PFU_NoUV_. Values are reported as mean +/− SD from n=3 experiments. (c) Infection of human lung cell line, HCT-8 from irradiated and untreated hCoV-OC43. Green fluorescence indicates infected cells while blue fluorescence indicates DAPI stains of nuclei. Images were acquired with a 40x objective, with the scale bars at 50 μm.

We next investigated the integration of hCoV-OC43 in human lung host cells after exposure to 300 seconds of UVC sources. **Fig. 2C** shows the representative images of human lung cells HCT-8 with hCoV-OC43 illuminated at different UVC wavelengths. We assessed the human cell lines for expression of the viral spike protein and found that 277-nm UVC outperforms the other UVC wavelengths in inactivation of coronaviruses.

### Examining rates of nucleotide degradation under different UVC wavelengths

To understand if the inactivation efficacy comes from RNA damage, we performed quantitative RT-PCR to examine the copy number of hCoV-OC43 after UVC irradiation. We observed that the copy number of hCoV-OC43 to be unperturbed after 300 seconds of 222-nm far UVC irradiation, while 254-nm UV lamp exerted the largest decrease in copy number followed by 277-nm UVC LED (**Fig. 3A**).

**Figure 3.**
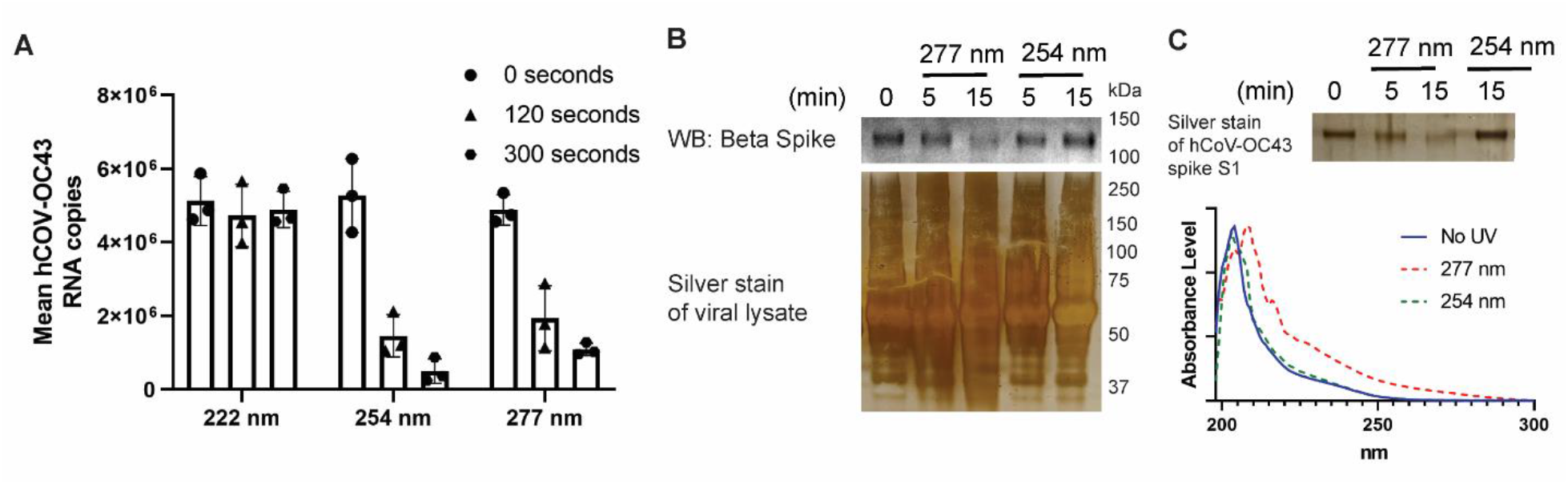
277nm UVC LED’s higher efficacy at inactivation of coronavirus is not due to UV-induced genomic damage and could be due to spike protein degradation. (a) Quantitative RT-PCR reveals that the copy number of hCoV-OC43 did not change due to 222-nm illumination, and decrease the fastest due to 254-nm UVC lamp. Values are reported as mean +/− SD from n=3 experiments. (b) Beta spike glycoprotein of hCoV-OC43 is found to diminish in intensity upon 15 minutes of 277-nm UVC LED illumination but not under 254-nm UVC lamp. Silver stain of viral lysate is provided to show the total loading on each lane. (c) Purified hCoV-OC43 spike S1 proteins is found to diminish in intensity upon 15 minutes of 277-nm UVC LED illumination but not under 254-nm UVC lamp. Changes in absorbance spectra of hCoV-OC43 spike S1 observed after 15 minutes of 277-nm UVC LED irradiation but not 254-nm UVC lamp.

### Photodegradation of hCoV-OC43 spike protein under 277-nm UVC LED

We next hypothesized that molecular components other than nucleic acids could be implicated, and the spike protein is an especially attractive target to pursue given that it facilitates viral transmission by binding to the host receptors(Shang et al., 2020). To this end, we subjected hCoV-OC43 to different duration of 254-nm and 277-nm UVC irradiation, and observed through western blot that the spike protein is degraded under 277-nm UVC LED and not under 254-nm UVC lamp. Silver staining of the viral lysate indicates the overall amount of protein loaded in each lane (**Fig. 3B**). To further confirm that spike protein is indeed degraded by 277-nm UVC LED, we performed UVC illumination on purified hCoV-OC43 spike proteins in vitro. Silver staining as depicted in **Fig. 3C** shows that hCoV-OC43 spike protein presents at a lower intensity under 277-nm UVC LED and not under 254-nm UV lamp. This is further corroborated by absorbance spectroscopy in **Fig. 3C** which demonstrates a shift in absorbance upon 277-nm UVC LED illumination but not 254-nm UV lamp.

### Photodegradation of SARS-CoV-2 spike protein under 277-nm UVC LED

First, we exposed SARS-CoV-2 spike protein S1 subunit to varied durations of 254-nm UVC lamp and 277-nm UVC LED and observed the reduction of the full-length spike protein band under 277-nm UVC LED and not under 254-nm UVC LED. This happens on both glycosylated and non-glycosylated forms of the spike protein (**Fig. 4A**). Absorbance spectroscopy (**Fig. 4B**) analysis showed the UV absorbance profile of 254-nm lit proteins to be relatively unchanged while there is an increase of absorbance profile in the 250-300 nm region for 277-nm lit proteins. Western blot (**Fig. 4C**) analysis further revealed a reduction in the SARS-CoV-2 spike S1 proteins under low loading of protein samples, but under higher loadings, aggregates of spike protein in the form of dimers and trimers could be seen in a dose-dependent manner.

**Figure 4.**
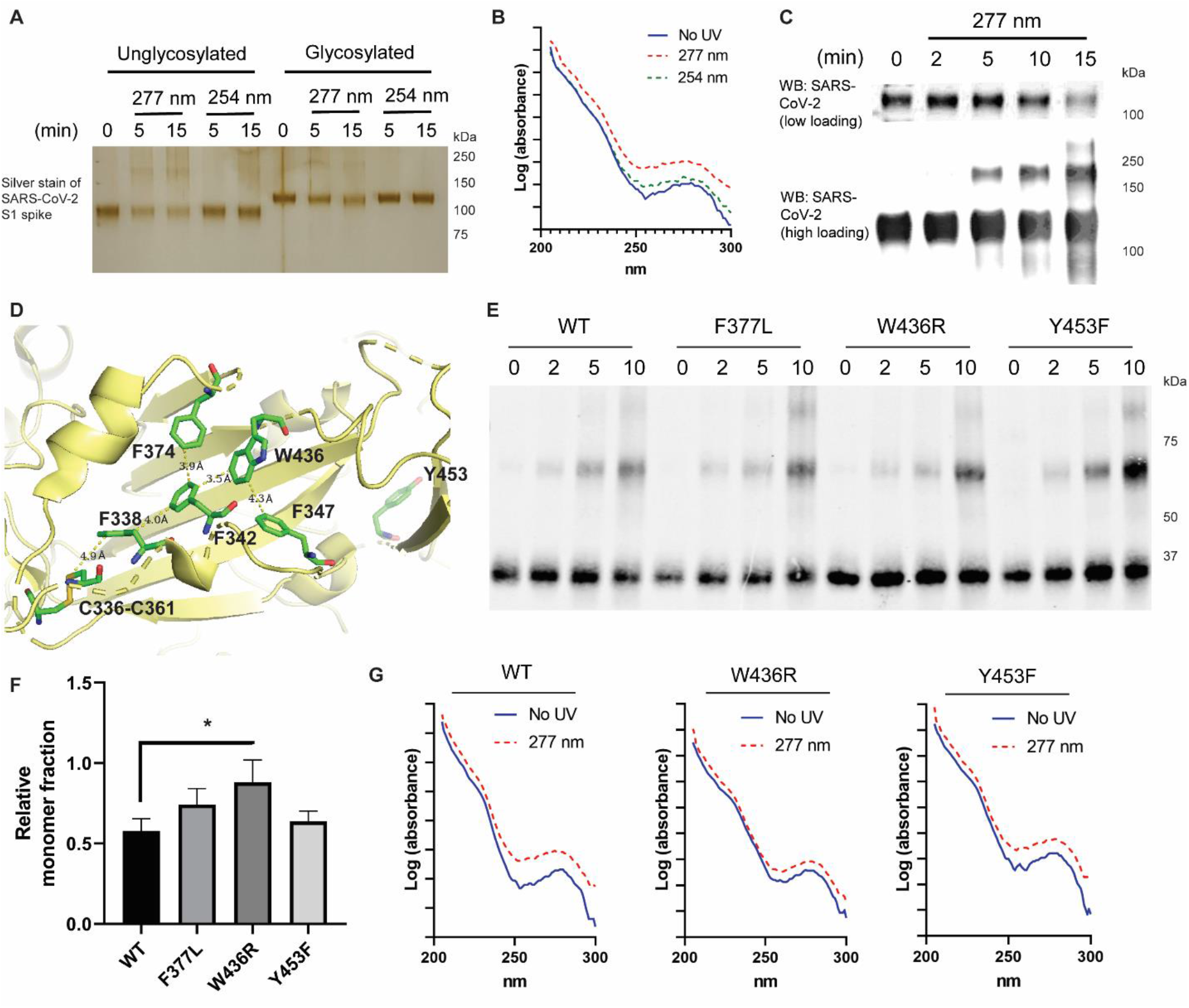
Photodegradation of SARS-CoV-2 spike protein under 277nm UVC LED. (a) Silver stains of un-glycosylated and glycosylated SARS-CoV-2 spike S1 protein under different UVC treatment. (b) Changes in absorbance spectra of SARS-CoV-2 spike S1 observed after 15 minutes of 277-nm UVC LED irradiation but not 254-nm UVC lamp. (c) Western blot analysis of SARS-CoV-2 spike S1 protein revealed that while the protein level of SARS-CoV-2 decreases under 277-nm UVC LED, higher aggregates could be observed upon higher UVC dose. (d) SARS-CoV2 spike protein structure (PDB: 6VXX). Key residues centered around W436 that could potentially act as an antenna for 277nm UVC absorption. Y453 is depicted in the background while F377 is adjacent to F374 (not highlighted in the schematic). (e) Western blot analysis of SARS-CoV-2 spike S1 RBD proteins revealed the differential rate of aggregation and degradation amongst the different mutants. (f) Quantification of relative monomer fraction for wild type and mutant RBD proteins. The fraction is calculated as a function of (intensity at 35kDa) / (overall intensity across the whole lane). (g) Absorbance spectra of the RBD proteins reveal little changes in 250-300 nm UVC absorbance for W436R compared to wild type and Y453F mutant, indicating potentially that W436R spike protein is less susceptible to 277nm UVC LED treatment.

In the search for potential mechanisms that drive the absorption of 277-nm wavelengths and degradation / aggregation of the proteins, we studied the structure of the SARS-CoV-2 S glycoprotein (PDB: 6VXX) (Walls et al., 2020) and looked for aromatic amino acids in proximity to disulfide bonds as possible active regions upon 277-nm UVC irradiation. In particular, tryptophan has high molar absorptivity at the 280-nm wavelength and has been known to mediate energy transfer and neighboring disulfide bond breakage (Beaven and Holiday, 1952; Chan et al., 2006; Kerwin and Remmele, 2007; Wu et al., 2008). We identified the Trp 436 as a key antenna of 277-nm absorption, and conducted studies on W436R receptor binding domain (RBD) mutant alongside wild type, F377L and Y453F mutants, that are in close proximity but unlinked to the W436-C336-C361 transfer chain (**Fig. 4D**). We illuminated the RBD samples with 0, 2, 5, 10 minutes of 277-nm UVC LED and studied the rate of oligomerization for each sample by probing with SARS-CoV-2 Spike antibody (**Fig. 4E**). It is observed that W436R mutant has a lower rate of aggregation compared to the other species and this is quantified by comparing the intensity of the monomer fraction (at 35kDa) with respect to the rest of the lane for samples illuminated with 5 minutes of 277-nm UVC LED (**Fig. 4F**). Absorbance spectroscopy further verified that after 10 minutes of UVC LED illumination, the W436R mutant did not exhibit as significant changes in absorbance compared to the wild-type and Y453F mutant.

## DISCUSSION

In this study, we focused on human coronaviruses hCoV-OC43 and hCoV-229E, which belong to the genus beta-coronavirus and alpha-coronavirus respectively. In particular, hCoV-OC43 could be considered as a surrogate for SARS-CoV-2 and the conclusions drawn here on UVC efficacy can thus be extrapolated to SARS-CoV-2. On the other hand, hCoV-229E resembles the viruses that causes the common cold. The viral efficacy tests performed here not only targeted the current COVID-19 pandemic, but also applies to future coronaviral pandemics in general.

While many studies have individually tested the 222-nm far UVC lamp, 254-nm UVC lamp and broad ranges of UVC LEDs for their efficacy towards viruses, this is the first study to report the mechanisms through which viral inactivation occurs. In summary, we find that 277-nm UVC LED outperforms the other UVC wavelengths in inactivation of human coronaviruses, and this could be aided by the contribution from spike protein degradation with absorption of UV wavelengths at Trp 436. We also find that 222-nm UVC LED does not affect the genomic material of beta-coronavirus, an observation that is congruent with a previous report (Kitagawa et al, 2020).

It has been widely believed that the efficacy of UV germicidal irradiation (UVGI) is dependent largely on the absorption by the target nucleic acids. While the mechanism holds significant merit, it is important to examine the other molecular mechanisms at which UV tools could exert their germicidal properties. It is thus important to consider the viral components individually as we characterize the multitude of UVGI solutions available to combat the current and future pandemics.

## LIMITATIONS OF STUDY

Due to the lack of access to BSL-3 facilities, we were unable to perform the viral infectivity tests on SARS-CoV-2 or the relevant variants. However, this limitation is mitigated by our studies on the beta coronavirus hCoV-OC43, which provides a close approximation towards SARS-CoV-2 with the relevant structures of the spike proteins being relatively similar.

## ASSOCIATED CONTENT

### Supporting Information

The following files are available free of charge.

**Figure S1.** Wavelength spectrum of different UVC light sources

**Figure S2.** Schematic diagram of UVC enclosure

**Table S1.** Two-way ANOVA results for hCoV-OC43 infectivity curves

**Table S2.** Two-way ANOVA results for hCoV-229e infectivity curves

### Author Contributions

The manuscript was written through contributions of all authors. Here entails the list of contributions made by each author:-Conceptualization: QO, JWRT, WH; Methodology: QO, JWRT; Investigation: QO, JWRT, JDC, EW, WW; Funding acquisition: QO, JWRT; Writing – original draft: QO, WH; Writing – review & editing: JWRT, JDC, EW, WW. All authors have given approval to the final version of the manuscript.

### Funding Sources

We would like to thank our funding sources:-National Research Foundation grant NRF2020NRF-CG002-035 (awarded to QO, JWRT) and Accelerate Technologies Gap-funded project GAP/2020/00392 (awarded to JWRT, QO).

## ACKNOWLEDGMENT

We thank Dr. Edward George Robins, Dr. Yaw-Sing Tan, Dr. Janarthanan Krishnamoorthy and Ms. Haitong Mao for proofreading of the manuscript; we thank Prof. Lisa Ng, Prof. Xianjun Loh, Dr. Yuanjie Liu, Ms. Siew Lan Lim and Dr. Sun-Yee Kim for providing technical advice and help.

## MATERIALS AND METHODS

### KEY RESOURCES TABLE

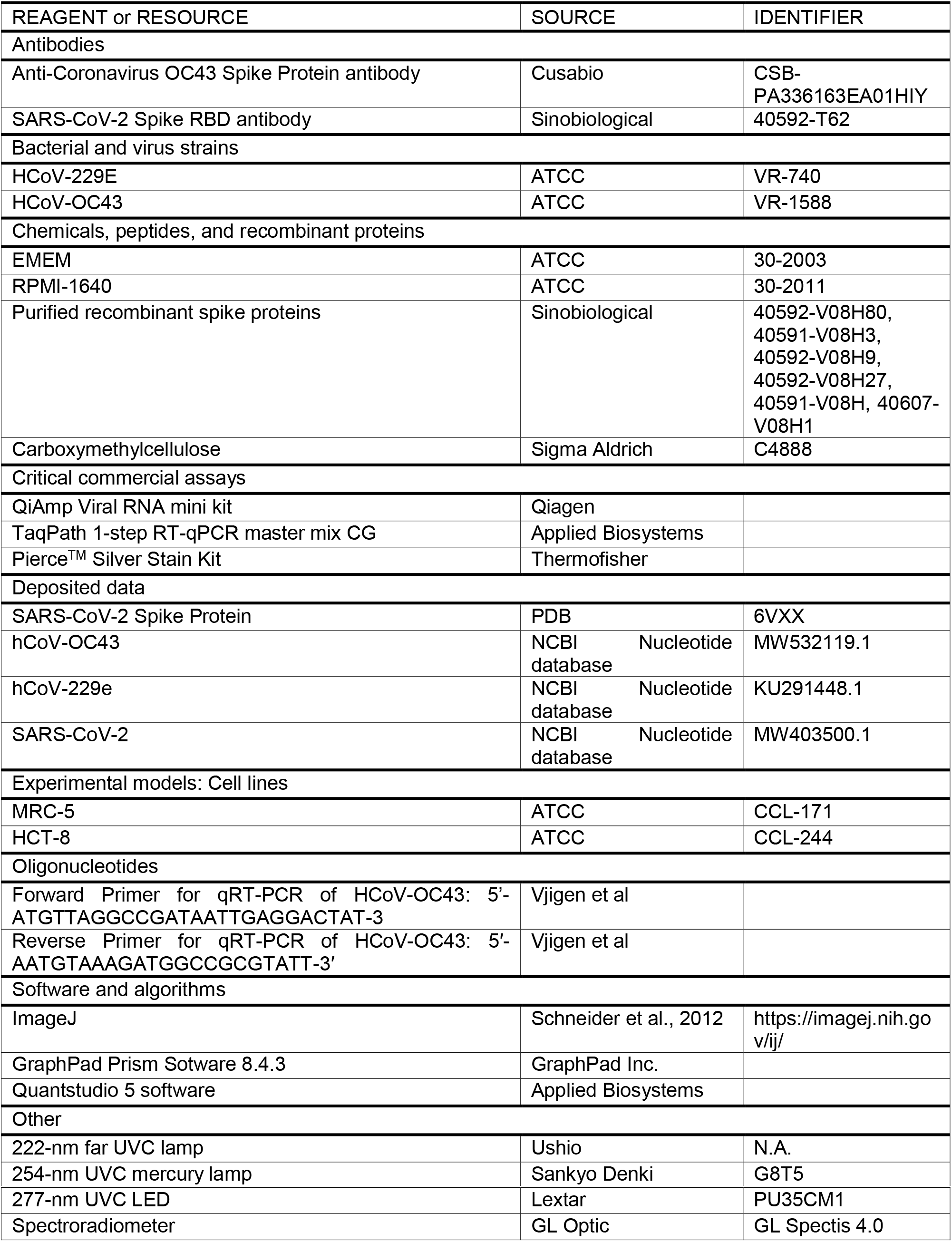

#### Viral strains and viral propagation

HCoV-229E (ATCC VR-740) and hCoV-OC43 (ATCC VR-1558) were propagated in human lung fibroblasts MRC-5 (ATCC CCL-171) and colon adenocarcinoma cells HCT-8 (ATCC CCL-244) respectively (all from ATCC, Manassas, VA). The MRC-5 fibroblasts were grown in EMEM (ATCC 30-2003) supplemented with 10% Fetal Bovine Serum (FBS), 100 U/ml penicillin and 100 μg/ml streptomycin (Sigma-Aldrich, St. Louis, MO). The HCT-8 epithelial cells were cultured in RPMI-1640 supplemented with 10% horse serum, 100 U/ml penicillin and 100 μg/ml streptomycin. The virus infection medium is made up of EMEM or RPMI-1640 with 2% FBS or horse serum for hCoV-229E and hCoV-OC43 respectively.

#### UV sources and irradiance measurement

To understand the effect of UVC wavelength on human coronaviruses, three different UVC light sources – 222-nm far UVC lamp (Ushio), 254-nm UVC mercury lamp (Sankyo Denki G8T5) and 277-nm UVC LED (Lextar PU35CM1) were used in this study. These UVC light sources were measured using a calibrated spectroradiometer (GL Spectis 4.0) with an absolute measurement uncertainty of less than 6%. To provide a comparative UVGI efficacy study between these UVC light sources, the radiant intensity of the far UVC and mercury lamp is measured at different distances while the UVC LEDs are driven at different constant drive currents to obtain a common UV intensity of 73 μW/cm^2^. The UVC LED, with a beam angle of 120°, is assembled into a 5 x 5 array at a working distance of 12 cm to ensure uniform UV intensity across the surface of the petri dish. The relative wavelength spectra of the light sources are shown in **Fig. S1**. Based on the UV intensity readings, an enclosure is fabricated for each light source and the radiant intensity of the enclosure is further validated with the spectroradiometer as shown in **Fig. S2**.

#### Viral infectivity experiments

The virus was propagated as previously described and stored in virus infection medium at 10^8^ PFU/ml. For each irradiation, 100 μl of virus suspension was placed on a 3-cm petri dish. After each irradiation, the virus was subjected to serial dilution of 5 times, and 50 μl of each condition is diluted with 450 μl of virus infection medium into 24-hour old and 80-90 % confluent MRC-5 or HCT-8 cells in 6-well plates. The cells were incubated with the virus for 1 hour in a humidified incubator with 5% CO_2_ before addition of a liquid overlay medium, 3% carboxymethylcellulose, is applied to the cells to restrict virus growth to the originally infected loci of cells. The cells are incubated at 37C in a 5% CO_2_ incubator for 3 days before the liquid overlay medium is aspirated and fixation with 4% paraformaldehyde is performed at room temperature for an hour. Staining with 0.5% crystal violet is then conducted and the plaques are then quantified.

#### Immunofluorescence experiments

To assess whether 5 minutes of various UVC illumination reduces the number of infected cells, immunostaining was performed to detect the presence of OC43 viral particles in the host human cells. Briefly, 2 x 10^5^ HCT-8 cells were plated in each petri dish one day before the experiment. The viral suspension after their respective treatments was overlaid on the monolayer of host cells. After one hour of incubation, the cells were washed with PBS and incubated for two days in fresh medium. The cells were then fixed with 4% paraformaldehyde at room temperature for 15 minutes and washed with PBS before being labelled with anti-Coronavirus OC43 Spike Protein antibody (CusaBio Technology LLC, Houston, TX, USA) 1:500 in PBS containing 2 % bovine serum albumin (BSA) and 0.1 % TBS-T. Cells were then washed with PBS and labelled with goat anti-rabbit Alexa Fluor-488 (Life Technologies, Grand Island, NY) in PBS containing 2 % BSA at room temperature for an hour with gentle shaking. Following washing with PBS, the cells were stained with DAPI and observed with the 40x objective of Nikon Ti-2 TIRF microscope.

#### Reverse Transcriptase Experiments

Beta-coronavirus RNA was extracted from the viral samples on each petri dish using the QiAmp Viral RNA mini kit (Qiagen) following the manufacturer’s instructions. 5 μl of each sample was used for qRT-PCR analysis utilizing the TaqPath 1-step RT-qPCR master mix CG (Applied Biosystems) and the primer set spanning a target region of 68bp for HCoV-OC43 as reported (20). The forward primer is 5’-ATGTTAGGCCGATAATTGAGGACTAT-3 and the reverse primer is 5’-AATGTAAAGATGGCCGCGTATT-3’. The analysis was then performed using the Quantstudio 5 software.

#### Protein degradation experiments

The antibodies that were utilized for western blots in this paper includes anti-Coronavirus OC43 Spike Protein antibody (CusaBio Technology LLC, Houston, TX, USA) and SARS-CoV-2 Spike RBD antibody (SinoBiological). All coronavirus purified spike proteins were obtained from SinoBiological. All samples for western blot were lysed in 2x Lamelli buffer and incubated in room temperature for 20 minutes before loading. Approximately 200 ng of protein sample is loaded for silver staining experiments and 1ug of protein sample is loaded for western blot experiments. The lysates were then subjected to SDS gel electrophoresis. For silver staining, we utilized the Pierce™ Silver Stain Kit (Thermofisher). For western blot, the samples were transferred to nitrocellulose membranes using iBlot2 (Life Technologies), blocked with 5% BSA in TBST and incubated with primary antibodies in 5% BSA. Membranes were then incubated with rabbit-IRDye 800 CW secondary antibodies and imaged on an Odyssey CLx (LI-COR). Absorption spectroscopy was performed on Nanodrop (Thermofisher).

#### Quantification and Statistical Analyses

The number of replicates (n) are indicated in the respective figure legends. For all statistical tests, significance was measured against p < 0.05. For comparisons of infectivity curves, 2-way analyses of variance (ANOVA) were carried out with the different UVC wavelengths and exposure time as the factors. All statistical analyses were performed with the GraphPad Prism Sotware 8.4.3 (GraphPad Software Inc., La Jolla, CA, USA) and all values are expressed as means +/− standard deviation.

#### Protein structure studies

The Desktop PyMOL 2.4 was used to visualize the protein structure of SARS-CoV-2 S glycoprotein (Protein Data Bank 6VXX) and calculate the distances between each indicated residue.

