## Supplementary Info for "Irradiation of UVC LED at 277 nm inactivates coronaviruses by photodegradation of spike protein"

**Supplementary Figures**

**Figure S1.** Wavelength spectrum of different UVC light sources

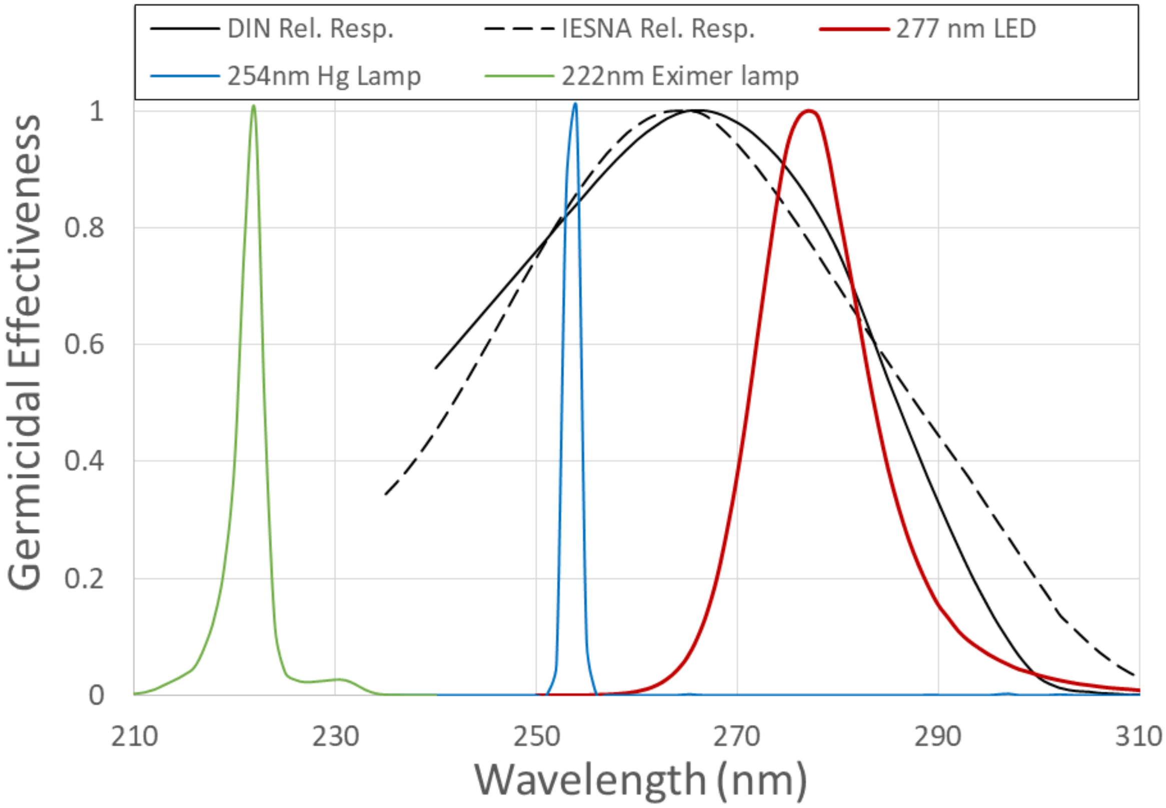

**Figure S2.** Schematic diagram of UVC enclosure

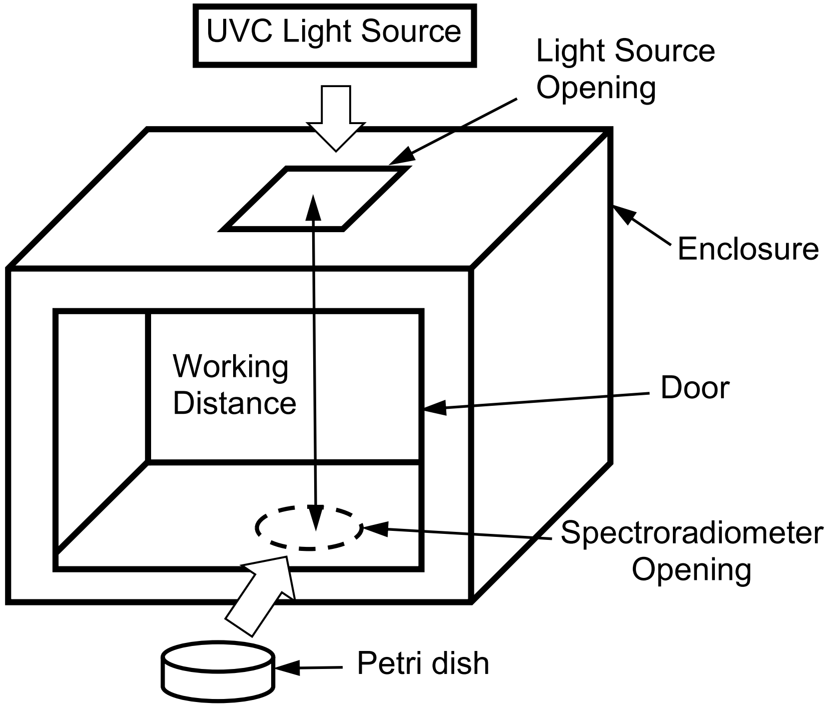

**Table S1.** Two-way ANOVA results for hCoV-OC43 infectivity curves.

| Source of Variation | % of total variation | P value | P value summary | Significant? | Geisser-Greenhouse's epsilon |
|---|---|---|---|---|---|
| Row Factor[a] x Column Factor[b] | 9.183 | <0.0001 | **** | Yes | |
| Row Factor | 85.96 | <0.0001 | **** | Yes | 0.4027 |
| Column Factor | 4.361 | 0.0001 | *** | Yes | |
| Subject | 0.2266 | 0.0628 | ns | No | |

| ANOVA table | SS | DF | MS | F (DFn, DFd) | P value |
|---|---|---|---|---|---|
| Row Factor x Column Factor | 6035 | 6 | 1006 | F (6, 18) = 100.7 | P<0.0001 |
| Row Factor | 56488 | 3 | 18829 | F (1.208, 7.249) = 1884 | P<0.0001 |
| Column Factor | 2866 | 2 | 1433 | F (2, 6) = 57.75 | P=0.0001 |
| Subject | 148.9 | 6 | 24.81 | F (6, 18) = 2.483 | P=0.0628 |
| Residual | 179.9 | 18 | 9.992 | | |

| Data summary | |
|---|---|
| Number of columns (Column Factor) | 3 |
| Number of rows (Row Factor) | 4 |
| Number of subjects (Subject) | 9 |

[a] Row factor refers to different timings of UVC exposure.

[b] Column factor refers to the different UVC wavelengths utilized.

**Table S2.** Two-way ANOVA results for hCoV-229e infectivity curves.

| Source of Variation | % of total variation | P value | P value summary | Significant? | Geisser-Greenhouse's epsilon |
|---|---|---|---|---|---|
| Row Factor x Column Factor | 6.090 | <0.0001 | **** | Yes | |
| Row Factor | 87.85 | <0.0001 | **** | Yes | 0.3988 |
| Column Factor | 5.446 | <0.0001 | **** | Yes | |
| Subject | 0.07287 | 0.8687 | ns | No | |

| ANOVA table | SS | DF | MS | F (DFn, DFd) | P value |
|---|---|---|---|---|---|
| Row Factor x Column Factor | 3633 | 6 | 605.5 | F (6, 18) = 33.51 | P<0.0001 |
| Row Factor | 52401 | 3 | 17467 | F (1.196, 7.178) = 966.6 | P<0.0001 |
| Column Factor | 3249 | 2 | 1624 | F (2, 6) = 224.2 | P<0.0001 |
| Subject | 43.47 | 6 | 7.245 | F (6, 18) = 0.4009 | P=0.8687 |
| Residual | 325.3 | 18 | 18.07 | | |

| Data summary | |
|---|---|
| Number of columns (Column Factor) | 3 |
| Number of rows (Row Factor) | 4 |
| Number of subjects (Subject) | 9 |
| Number of missing values | 0 |

[a] Row factor refers to different timings of UVC exposure.

[b] Column factor refers to the different UVC wavelengths utilized.
